# The date palm (*Phoenix dactylifera* L.) in Siwa oasis (Egypt): How ethnographic, morphometric, and genetic analyses together explain the local agrobiodiversity

**DOI:** 10.1101/122846

**Authors:** Vincent Battesti, Muriel Gros-Balthazard, Clémence Ogéron, Sarah Ivorra, Jean-Frédéric Terral, Claire Newton

## Abstract

The agrobiodiversity of the Siwa oasis (in Egypt), located at the crossroads of ancient Trans-Saharan routes, is evaluated in this article focusing on the date palm (*Phoenix dactylifera* L.), the oasis “ecosystem engineer”. This assessment confronts different ontologies: diversity as expressed and maintained by the folk categorization system of the Siwa inhabitants (through the results of an ethnographical analysis) and diversity described by genetic sciences and a morphometric tool based on the size and geometry of the seeds. This work is also an opportunity to evaluate this tool intended for archaeobotany. Beyond a simple instrumentalization of one discipline by another, this study offers a space of mutual enrichment: on the relative importance of the feral and cultivated date palms, the local relevance of the concept of “cultivar” and the confirmation of the existence of “ethnovarieties”.

## Introduction

Agrobiodiversity or agricultural biological diversity refers to the variability of animal, plant and microbial organisms maintained by human societies. Through agricultural practices carried on generations, human societies have a direct impact on their environment, partly shaping it by creating and organizing biodiversity. Agrobiodiversity has been and still is also a tool to cope with current and future challenges such as shifting environments (including the emergence of new diseases), climate change, and changing socioeconomic demands. The assessment and the conservation of agricultural diversity is therefore of major interest (Gepts et al. 2012). However, the appreciation of agrobiodiversity is difficult as it is intrinsically related to cultural practices and local knowledge. A sole biological approach is always inefficient to correctly assess the agrobiodiversity.

The date palm (*Phoenix dactylifera* L.) has been grown for millennia around the Persian Gulf and in North Africa (Barrow 1998). It was domesticated from wild populations of the same species but none is yet clearly identified (Pintaud et al. 2010), although some Omani populations have recently been proposed (Terral et al. 2012, Gros-Balthazard et al. 2016). Until recently, it was thought it had been domesticated in the Persian Gulf but recent genetic data suggests a second possible domestication or diversification center in Africa (Hazzouri et al. 2015, Mathew et al. 2015, Zehdi-Azouzi et al. 2015). The date palm is the cornerstone of ingenious oasis agrosystems, oases being artificial structures invented, sometimes very anciently, and maintained by local people (Battesti 2005).

Numerous cultivars have been selected by local populations, especially in oases, to meet the local and regional needs and local pedoclimatic conditions. It is quite impossible to have an accurate number of these hundreds of cultivars currently grown. A general survey indicates that more than 3,000 cultivars probably exist (Zaid & Arias-Jiménez 1999) and probably more than 200 in Tunisia only (Rhouma 1994, 2005). However, the notion of cultivar is complicated. It is assumed to be a collection of clones that is the association of a name and single genotype (reproduced vegetatively/asexually). For instance, all Deglet Noor date palms are supposed to derive from a unique plant through vegetative propagation of offshoots. Nevertheless, genetic data demonstrated that cultivars are not necessarily 100% identical (for instance, Medjool from Morocco; Elhoumaizi et al. 2006). Additionally, some cases of homonymy (same name used in different localities for different genotypes) have already been noted in the literature, in Egypt (El-Assar et al. 2005: 606), in Libya (Racchi et al. 2013) and elsewhere (Khanamm et al. 2012: 1240). Lastly, intra-cultivar genetic variability might exist due to somatic mutations (Devanand & Chao 2003). Therefore, the notion of cultivar, resulting from social (a name) and agricultural work (a genotype), still remains debatable.

In Siwa oasis (Egypt), the notion of cultivar is not expressed in the *jlan en Isiwan* (the local Amazigh language, a variety of Berber language), even if the notion of cultivar seems there to be an agronomic reality. According to our ethnobotanical results (Battesti 2013), the agrobiodiversity of date palms in Siwa seems organized according to the notion of *eškel* (pl. *eškolĩ*) which refers to a “form” (meaning of *šakl*, the Arabic root of this “berberized” word) of date palm associated to a name, and this equates either with a cultivar, an ethnovariety, a landrace or a category of date palms. For farmers, to focus on numbers to determine the agrobiodiversity is meaningless: “but there are hundreds of [forms of] different date palms! Most of the *úšik* [see further] do not have a name!”. In Siwa oasis, in addition, it has been shown that a single cultivar may bear different names and that a diversity of seedlings with their own denomination grows in the gardens (Battesti 2013).

The oasis of Siwa is located in the midst of the Libyan Desert, Egyptian territory, 300 km south of the Mediterranean coast and the closest city Marsa Matruh, and about 70 km east of the Libyan border (Figure 1). Siwa is the name of the small region, a set of palm groves and settlements, but also nowadays the name of its main “town”. The area is situated in a depression about 50 km in length and 10 km wide, a dozen meters below sea level. Siwa oasis is often described in the literature as a remote and forgotten place (but a never-to-be-forgotten place for its visitors). Nonetheless, Siwa has not been “forgotten” by everyone: first its inhabitants, but also all the networks Siwa is linked to. Siwa was for times immemorial (*stricto sensu*) an inevitable node in the East-West and North-South Saharan routes networks (Battesti in press). All land communication and land transport of goods and living material (such as cultivated plants) between North Africa and the Nile Valley/Middle East have to go through this “oasian port”, Siwa, which changed its name in the course of centuries and millennia (Sekhet-Ȧmit, Santar, Amon, Jupiter-Ammon, Marmaricus Hammonn, Hammoniacus Nomos, Santariyya, Siwa). Dates from Libyan oases were exported to the Greek world through Siwa at least from the fifth century BC (review in Leclant 1950) and dates from Siwa itself were particularly famous in the first century AD (Plinus, Natural History, XIII, 111, in Leclant 1950: 248).

**Figure 1.**
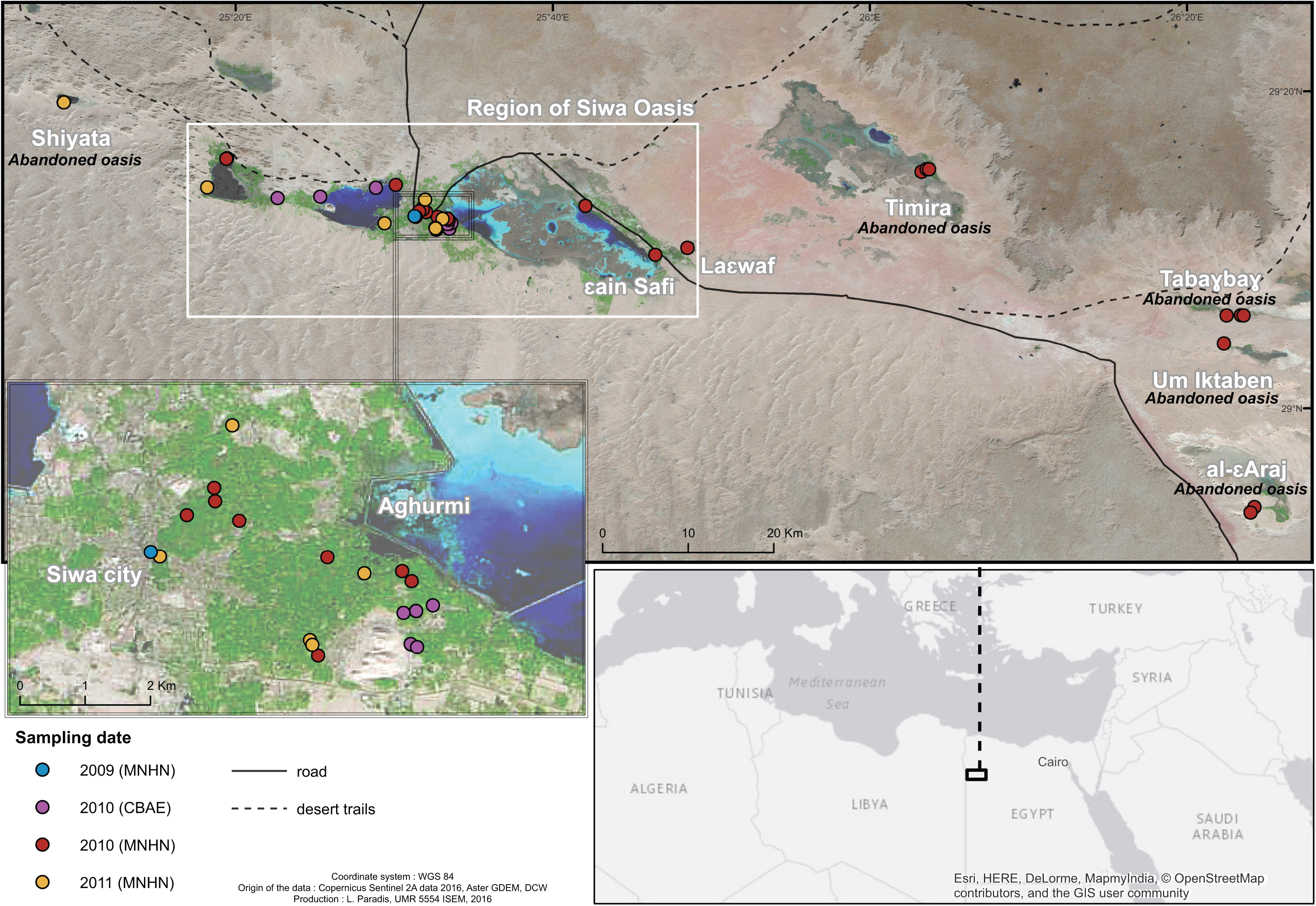
Localization of the main geographic locations of the present study accessions.

In Siwa, the date palm constitutes the main cash crop of the oasis (Battesti 2013). The fame of the date palm of Siwa dates back to Antiquity (Siwa’s name in ancient Egyptian, Sekhet-Ȧmit, means “field of the palm trees”). But until now no accurate idea of the agrobiodiversity of these date palms has been produced despite more than 60 references in all languages published over the last two centuries mentioning varieties of dates from Siwa, but often confusing local names of categories of dates with varieties and cultivars (or so-called varieties and cultivars), and despite some recent published surveys of its genetic diversity (Adawy et al. 2005, El-Assar et al. 2005, Hussein et al. 2005, Rizk & Rashed 2005, Abou Gabal et al. 2006, Soliman et al. 2007, Abd El-Azeem et al. 2011). This paper focuses on the agrobiodiversity of date palms in Siwa. Our ethnoecological survey (Battesti 2013) explained this local difficulty to draw up the date palm biodiversity list of Siwa and offered a possible overall view, still to be confirmed in its hypothesis by genetic analyses (see Figure S1). The ethnographic study revealed that the farming practices are probably different from what they are supposed to be. It is usually considered that oasis farmers in general asexually reproduce date palm cultivars, that the latter are collections of clones (each collection with a given name), and that reproduction only occurs through the production of offshoots by the palm trees: “As a vegetatively propagated perennial fruit tree, the date palm is unique in that it is composed of genetically discrete clones representing highly heterozygous cultivars without the benefits of a dynamic mutation-recombination system.” (Jaradat 2015: 20) It is also often assumed that seedlings date palms are only accidents from seeds and ignored by local oasis farmers (Battesti 2005).

This study is the first to offer an analytical list of the date palm agrobiodiversity of Siwa Oasis. What we know so far is this (Battesti 2013): we have about fifteen different named types in Siwa only, ignoring the different intermediary names (Figure S1). To summarize drastically a complex situation, we can say that the distinction is locally first made between two elite cultivars, *Ṣaεidi* and *alkak*, the two main cultivated, harvested and exported dates in Siwa. Both keep and travel well thanks to their low moisture content and high sugar content; *Ṣaεidi* is especially exported to cities for confectionery and *alkak* is consumed by the Bedouins transporting the *Ṣaεidi* from Siwa. In second place come the other alleged cultivars, locally appreciated but less common: *a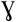zāl, alkak wen žemb, lekrawmet, ta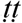agt, úšik niqbel, 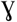rom a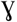zāl*, and *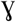rom Ṣaεid*. In third place come some controversial named types: Siwa inhabitants all agree on the fact these date palms are vegetatively reproduced (cloned, as the “alleged true cultivars”) but they do not all agree on the possibility that a seedling can give by chance one of these named types. These named types are candidates to be what has been called an ethnovariety. Our current definition of an ethnovariety for the date palms is: “set of similar (according to local standards) lines of clones reproduced by vegetative offshoot under the same local name” (Battesti 2013). An ethnovariety differs from a landrace. An ethnovariety is always reproduced vegetatively, but might benefit from a seedling to create a new line of clones. According to Pintaud (2010: 109), a landrace of date palms is a group of date palms partially clonal or fully propagated by seed, but by consanguinity retaining a stable set of morphological traits. This local observation can perhaps explain the genetic differences within a so-called single line of clones of a cultivar, as reported in several genetic studies elsewhere (which usually put forward the hypothesis of somatic mutations). Those controversial named types in Siwa are: *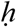alu en 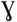anem, amenzu, úsik ama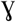zuz, úsik ezzuwa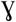* (or *zuwa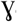*), *úšik azzuga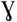, tažubart*. To that, we must add two most probable ethnovarieties: *kaεibī* and *úšik nekwayes*. Finally, to complete the picture, we should mention the date palm seedlings. Actually, as subspontaneous, they are not really part of the date palm agrobiodiversity, but its result, and are nonetheless an important gene pool. Usually, seedlings are discarded from the garden, but some nevertheless grow, are not uprooted by the famers, may even be pollinated and tended because, against all expectations, their fruit are good, or a good fodder, or anything else. Those garden seedlings are called *úšik*. We call them in this paper *úšik* #1. The seedlings that grow untended outside the garden in the collective part of the palm grove or in its immediate periphery are also locally called *úšik*. We call them in this paper *úšik* #2. Finally, date palms from abandoned oases in the desert nearby Siwa are today mainly propagated by seed (plus their natural offshoots), They grow spontaneously without human intervention even if the date palms were maybe formerly cultivated. As subspontaneous, they are “feral” date palms; the population of *úšik* date palms in Siwa, outside the gardens (*úšik* #2), may also be considered as feral.

The aim of this study is to verify the validity of these statements in the oasis of Siwa. For that purpose, we used a combination of three different approaches: one approach specifically from the social sciences (ethnographical/ethnobotanical analysis) and two approaches specifically from the biological sciences (seed morphometrics and genetic analyses). An ethnographical approach makes accurate morphometric and genetic interpretation possible, first by collecting correctly the local given names of date palms, secondly by researching the use and qualification of local dates and finally by reviewing the local naming and classification of date palms from scratch. Ultimately, the objective is to obtain a better assessment of the local folk categorization. Genetic analyses will allow to accurately verify the relationships of the samples using microsatellite loci as already employed in Bodian et al. (2014) or Elhoumaizi et al. (2006). Furthermore, we performed seed morphometric analyses. This approach has extensively been used in date palms for characterizing past and present agrobiodiversity and hypothesizing the existence of wild populations (Terral et al. 2012, Gros-Balthazard et al. 2016). Indeed, the size and shape of seeds are indicators of the status of the plants (wild, feral or cultivated) and vary between cultivars (Terral et al. 2012, Gros-Balthazard et al. 2016). The main advantage is that it can be applied to archaeological samples. Indeed, archaeobotanists need tools to trace back the evolution of the date palm and its biodiversity, especially in the context of excavations under arid climates, where DNA is rarely preserved well enough for analysis. The seed and its shape *a contrario* are well preserved. The objective of this study was to validate this tool for the date palm thanks to a double analysis, morphometric and genetic, on contemporary samples of date palms from Siwa, informed by the ethnographical data. The examination of date seeds will first allow a morphological evaluation of the date palm diversity in Siwa (both cultivated and uncultivated). Additionally, seeds from multiple date palms belonging to a single true cultivar (i.e. clone) are expected to display identical shapes and we will thus assess if this is the case or not. The double analysis (morphometric and genetic) of samples from the same date palm offers the opportunity to blindly crosscheck the results and in return to test the local categorization of date palms collected by ethnographical fieldwork. The statistical procedures, and especially the interpretations of the statistical results, prompted the social and biological scientists to work together. The complementarity of their joint expertise was a pressing necessity to address a plant that mostly exists in a cultivated state. Its diversity is not only driven by human practices but has to be interpreted with local expertise and from a local perspective.

## Material & Methods

### Sample collection

A total of 62 date palm accessions were sampled for leaves and/or seeds in and around Siwa oasis (Table 1; Figure 1). None of our specimens came from experimental stations, but from *in situ* local (private) gardens in palm groves or their periphery. A palm grove is there as elsewhere not a mere palm forest, but a mosaic of private cultivated gardens, most of the time with mixed farming (Battesti 2005). We usually sampled about 40 seeds per specimen for morphometric analyses, along with a few leaflets per specimen for genetic analyses. Two specimens were collected in 2009 (seeds only). Then, two different drives took place during the Fall of 2010: their outcomes were 14 specimens for the first drive and 38 specimens for the second (seeds and leaflets), distributed respectively among six and 13 relevant given names. These drives were completed in the Fall of 2011 with 14 specimens (seeds and leaflets) distributed among six relevant given names.

**Table 1.** Date palm samples and the type they belong to. The status (feral, seedling or cultivated) is mentioned along with whether or not the sample was studied using seed morphometric and/or genetic.

| ID | Type | Status | Morphometric analysis | Genetic analysis |
| --- | --- | --- | --- | --- |
| 2048 | úšik #1 | garden seedling | yes | yes |
| 2049 | úšik #1 | garden seedling | yes | yes |
| 2051 | alkak | garden cultivated variety | yes | yes |
| 2052 | tattagt | garden cultivated variety | yes | yes |
| 2054 | úšik #2 | palm grove seedling | yes | yes |
| 2055 | aɣzāl | garden cultivated variety | yes | yes |
| 2056 | úšik #1 | garden seedling | yes | yes |
| 2058 | úšik ezzuway | garden cultivated variety | yes | yes |
| 2060 | úšik #1 | garden seedling | yes | yes |
| 2062 | úšik ezzuway | garden cultivated variety | yes | yes |
| 2063 | úšik #2 | palm grove seedling | yes | yes |
| 2065 | yrom aɣzāl | garden cultivated variety | yes | yes |
| 2397 | kaɛibī | garden cultivated variety | yes | no |
| 2398 | šaɛidi | garden cultivated variety | yes | yes |
| 2399 | šaɛidi | garden cultivated variety | yes | yes |
| 2400 | šaɛidi | garden cultivated variety | yes | yes |
| 2401 | šaɛidi | garden cultivated variety | yes | yes |
| 2402 | úšik #1 | garden seedling | yes | yes |
| 2403 | úšik #1 | garden seedling | no | yes |
| 2404 | šaɛidi | garden cultivated variety | yes | yes |
| 2405 | tattagt | garden cultivated variety | yes | yes |
| 2406 | úšik nekwayes | garden cultivated variety | yes | yes |
| 2407 | úšik #1 | garden seedling | no | yes |
| 2408 | alkak | garden cultivated variety | yes | no |
| 2409 | aɣzāl | garden cultivated variety | yes | yes |
| 2410 | úšik amayzuz | garden cultivated variety | yes | yes |
| 2411 | úšik #1 | garden seedling | yes | yes |
| 2412 | úšik ezzuway | garden cultivated variety | yes | yes |
| 2413 | *feral* | desert feral | no | yes |
| 2414 | *feral* | desert feral | yes | yes |
| 2415 | úšik #1 | garden seedling | no | yes |
| 2416 | alkak | garden cultivated variety | yes | yes |
| 2417 | alkak wen žemb | garden cultivated variety | yes | yes |
| 2418 | úšik #2 | palm grove seedling | yes | yes |
| 2419 | úšik ezzuway | garden cultivated variety | yes | yes |
| 2420 | *feral* | desert feral | no | yes |
| 2421 | *feral* | desert feral | yes | yes |
| 2422 | úšik niqbel | garden cultivated variety | yes | yes |
| 2423 | úšik niqbel | garden cultivated variety | yes | yes |
| 2424 | úšik #2 | palm grove seedling | yes | yes |
| 2425 | *feral* | desert feral | yes | yes |
| 2426 | *feral* | desert feral | yes | yes |
| 2427 | *feral* | desert feral | yes | yes |
| 2428 | *feral* | desert feral | yes | yes |
| **2429** | úšik #2 | palm grove seedling | no | yes |
| **2430** | *feral* | desert feral | no | yes |
| **2431** | *feral* | desert feral | yes | yes |
| **2432** | *feral* | desert feral | yes | yes |
| **2433** | *feral* | desert feral | yes | yes |
| **3289** | šaēidi | garden cultivated variety | yes | no |
| **3290** | šaēidi | garden cultivated variety | yes | no |
| **3291** | šaēidi | garden cultivated variety | yes | no |
| **3292** | šaēidi | garden cultivated variety | yes | no |
| **3293** | alkak | garden cultivated variety | yes | no |
| **3294** | ḡrom ayzāl | garden cultivated variety | yes | no |
| **3295** | šaēidi | garden cultivated variety | yes | no |
| **3296** | úšik niqbel | garden cultivated variety | yes | no |
| **3298** | šaēidi | garden cultivated variety | yes | no |
| **3299** | úšik #2 | palm grove seedling | yes | no |
| **3300** | úšik #2 | palm grove seedling | yes | no |
| **3301** | *feral* | desert feral | yes | no |
| **3302** | *feral* | desert feral | yes | no |

### Ethnobotanical study

The major material and methods of social anthropology discipline are time and fieldwork, well-equipped in observation and interview expertise, plus ethnobotanical knowledge: the social scientist (social anthropologist) undertook most of the *in situ* sampling drives. For a general survey of the method used to collect qualitative data such as local given names to the different kinds of date palms and to each of them separately, we refer to Battesti (2013). Data collection was backed up by years of work experience and long-term observations and interviews in the field with local farmers in Siwa oasis.

### Genetic analyses

Total cellular DNA was extracted from dried leaves of 47 samples, which were then genotyped using 17 microsatellite loci (Table 2) following the protocol of Zehdi-Azouzi et al. (2015). The structure of the genetic diversity in Siwa was assessed with a Principal Component Analysis (PCA) on gene frequencies using *dudi.pca* function (*ade4* package: Dray & Dufour 2007), R software, R Core Team (2015). A Neighborjoining (NJ) tree was constructed with Nei’s genetic distances (Nei et al. 1983) using *aboot* function in R (*poppr* package: Kamvar et al. 2014). The similarity between accessions from the same given names was calculated as the average percentage of loci that are not polymorphic. We expect that a collection of clones is 100% similar except in case of somatic mutations.

**Table 2.** Overview of microsatellite loci.

| Marker | Reference | missing data | % missing data | # alleles |
| --- | --- | --- | --- | --- |
| mPdCIR085 | Billotte et al. 2004 | 0 | 0.00 | 10 |
| mPdCIR078 | Billotte et al. 2004 | 1 | 1.06 | 11 |
| mPdCIR015 | Billotte et al. 2004 | 30 | 31.92 | 4 |
| mPdCIR016 | Billotte et al. 2004 | 1 | 1.06 | 5 |
| mPdCIR032 | Billotte et al. 2004 | 4 | 4.26 | 6 |
| mPdCIR035 | Billotte et al. 2004 | 0 | 0.00 | 3 |
| mPdCIR057 | Billotte et al. 2004 | 0 | 0.00 | 6 |
| mPdCIR025 | Billotte et al. 2004 | 2 | 2.13 | 8 |
| mPdCIR010 | Billotte et al. 2004 | 4 | 4.26 | 8 |
| mPdCIR063 | Billotte et al. 2004 | 4 | 4.26 | 6 |
| mPdCIR050 | Billotte et al. 2004 | 2 | 2.13 | 10 |
| PdAG1-ssr | Ludeña et al. 2011 | 1 | 1.06 | 12 |
| PdCUC3-ssr1 | Zehdi-Azouzi et al. 2015 | 2 | 2.13 | 1 |
| mPdIRD013 | Aberlenc-Bertossi et al. 2014 | 1 | 1.06 | 2 |
| mPdIRD033 | Aberlenc-Bertossi et al. 2014 | 2 | 2.13 | 4 |
| mPdIRD031 | Aberlenc-Bertossi et al. 2014 | 4 | 4.26 | 3 |
| mPdIRD040 | Aberlenc-Bertossi et al. 2014 | 1 | 1.06 | 5 |
| **Overall** |  | **3.53** | **3.69** | **6.12** |

**Table 3.** Average percentage of similarity calculated with genetic data within each named type of Siwa.

| Variety | # accessions | Percentage |
| --- | --- | --- |
| alkak | 2 | 91.18 |
| alkak wen žemb | 1 | / |
| ayzāl | 2 | 60.61 |
| *desert feral* | 12 | 17.65 |
| šaēidi | 5 | 96.97 |
| tattagt | 2 | 96.77 |
| úšik #1 | 8 | 18.18 |
| úšik #2 | 8 | 26.47 |
| úšik amayzuz | 1 | / |
| úšik ezzuway | 2 | 41.18 |
| úšik nekwayes | 1 | / |
| úšik niqbel | 2 | 65.63 |
| yrom ayzāl | 1 | / |

### Morphometric analysis of seeds

A total of 1100 seeds (20 per sample) from 55 accessions (Table 1) were analyzed using both traditional (size) and geometric (shape) morphometrics. The method associated with the quantification of seed size and shape has been described in Terral et al. (2012) and Gros-Balthazard et al. (2016). Briefly, seeds were photographed in both dorsal and lateral sides. Their length and width (cm) on the dorsal side (following Gros-Balthazard et al. 2016) were measured using ImageJ (Abràmoff et al. 2004). The shape of seeds was quantified using the Elliptic Fourier Transform (EFT) method (Kuhl & Giardina 1982) and implemented in R software (Momocs package, Bonhomme et al. 2014). To summarize, seed outline was extracted semi-automatically to obtain *x* and *y* coordinates of 64 points equally spaced along the outline of the seed. To avoid information redundancy and to standardize size and orientation, the coordinates were transformed using the EFT method leading to 32 harmonics, of which eight were retained (see Terral et al. 2012), with *A_n_* and *B_n_* related to x coordinate, and *C_n_* and *D_n_* related to *y* coordinate. The harmonic coefficients were used for subsequent statistical analyses in R.

The difference in seed size between the named types of Siwa was tested using Tukey’s test (function *HSD.test*, library *agricolae*). We appreciated the level of variability in both seed size and shape, according to the protocol developed in Gros-Balthazard et al. (2016). To summarize, we first performed a PCA on size or shape measurements and estimated the variation by calculating the size of the cloud valued as the mean distance between 10 seeds from a given group randomly sampled and the centroid of that given group. Of course, this method makes sense only if several accessions from the same “cultivar” are included. We expected a true cultivar to show a reduced seed size variation while heterogeneous groups should have seeds of different sizes and shapes, and therefore show a high seed size or shape variation. We further explored the structure of seed morphotype diversity using a Principal Component Analysis (PCA) performed using both discrete and outline measurements (function *dudi.pca*, package *ade4*). Additionally, we performed three Linear Discriminant Analyses (LDAs) on this full dataset. This aimed at assessing whether seed size and shape allow, on the one hand to discriminate uncultivated date palms (desert feral and palm grove seedlings) from cultivated date palms (named types and garden seedlings), then desert feral date palms from garden named types and from seedlings growing either in palm groves or gardens; and on the other hand to discriminate cultivars/named types from each other. We thus performed LDAs (*lda* function, package MASS) and used either factor “type” or “status” for the discrimination. To estimate the discriminant power of the LDAs, leave-one-out cross-validations were performed: posterior assignations were executed for each seed (option CV = T). The discriminating rate of each status or type was calculated as the percentage of positive allocation.

## Results

### Ethnobotanical study

The results of the ethnobotanical study were fully explained in Battesti 2013 and a summary is given in introduction of this paper and in Figure S1. Let us return briefly to this notion of feral in Siwa, because it is an important point to grasp the dynamics of agrobiodiversity. The term *úšik* is often part of a cultivar name, insisting on its former quality of *úšik*. This qualification emphasizes the origin of the genetic lineage as the outcome of a seed (such subspontaneous seedling date palms are said *úšik*): the farmer let such a female palm grow and then selected it, reproduced it and named it for its quality. The only “socialized” date palms in Siwa come from vegetative reproduction and are cultivated. A larger genetic stock however exists. First as *úšik* seedlings inside gardens (*úšik* #1 in this paper). This subspontaneous *úšik* of the cultivated areas has a counterpart in the communal spaces of the palm grove and its margins (*úšik* #2 in this paper) or in the abandoned palm groves (for centuries, not to say since Antiquity, probably the late Roman period) scattered in the Libyan desert around Siwa: *agzzo* pl. *igizzã*, in all likelihood also feral date palm seedlings, subspontaneous escaped from cultivation.

### Genetic analyses

The genetic analysis consisted in genotyping 47 accessions using 17 microsatellite loci (SSRs). The percentage of missing data is low (Table 2), indicating the efficiency of the microsatellite genotyping procedure. A total of 104 alleles were observed over the 17 SSRs, with an average of 6.12 per locus (Table 2).

#### Overall genetic diversity

The genetic structure of Siwan date palm diversity was inferred using both a NJ tree (Figure 2) and a PCA (Figure 3). In the PCA, the Principal Components (PCs) 1 and 2 explain the largest part of the diversity structure and almost the same amount (10.82% and 10.67% respectively). The PC1 is obviously stretched by *alkak* and *ta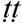agt*, the first is sweet and dry when the second is aqueous and soft (very estimated, but too aqueous to be preserved and to travel). PC2 clearly separates the two main cultivars from Siwa, namely *Ṣaεidi* and *alkak*. The distinction between these three named types (*alkak, Ṣaεidi, ta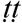agt*) is also evidenced in the tree (Figure 2) where they form distant clusters. PC2 is also drawn by the opposition between *úšik* (both #1 and #2) and desert feral date palms. PC1 and 2 together also separate desert feral from cultivated samples despite a little overlap. This distinction is also seen in the tree where all feral samples and some of the *úšik* #1 and #2 form a distinct clade (along with a few cultivated accessions, see below). The principal component 3 (Figure 3) does not show us more except that it separates *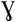rom a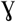zāl* and *ta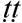agt* from the feral desert date palms, which was not the case with PC1 and 2 only.

**Figure 2.**
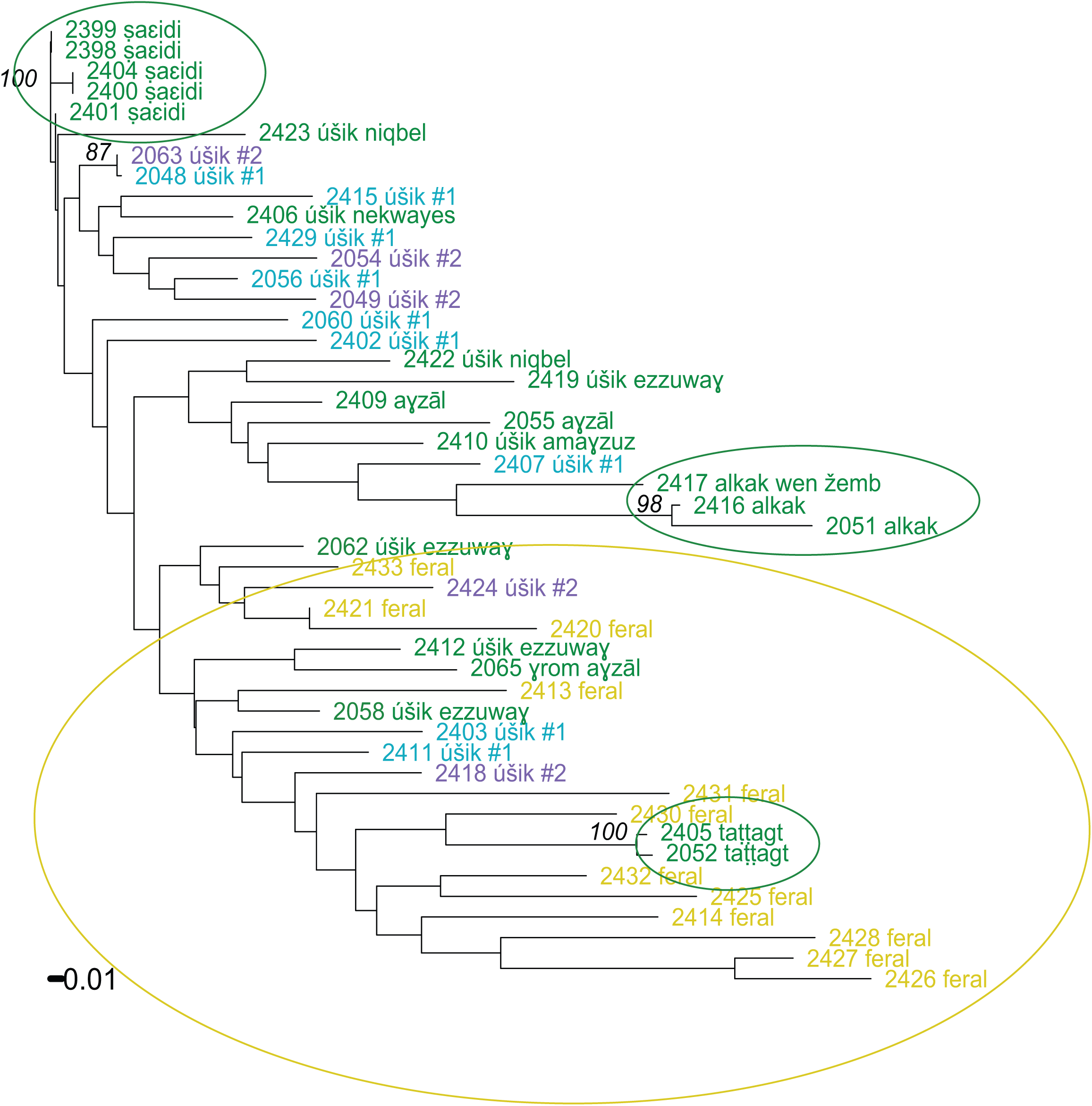
Neighbor-Joining (NJ) tree of Siwa date palms based on Nei 1983 genetic distances

**Figure 3.**
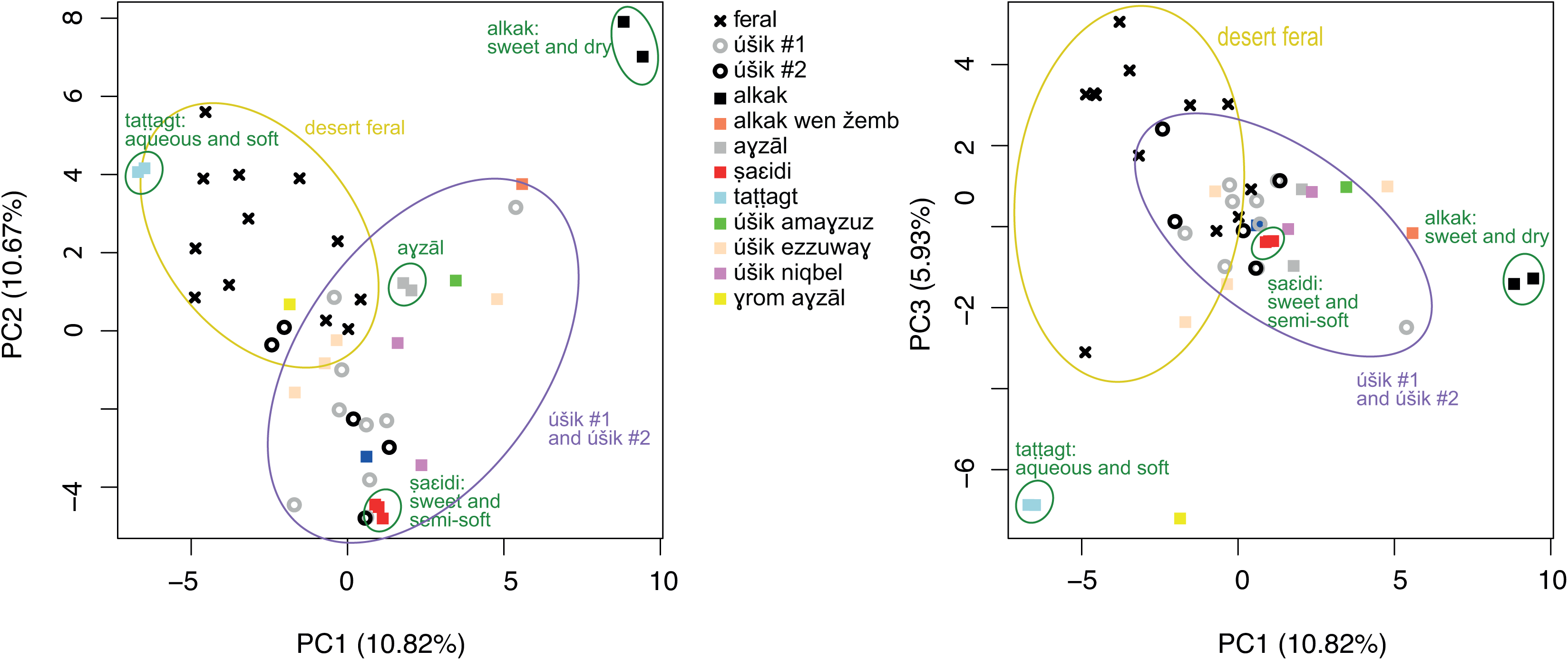
Principal Component Analysis (PCA) on 48 date palm accessions using SSRs data.

The whole two-dimension space shaped by PC1 and PC2 together seems shaped by the combination of two important sets of qualities: aqueous/sweet and soft/semi-soft/dry. The center of the graph, taken by the *a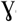zāl*, confirms this reading: a cultivar with semi-soft and sweet dates. Those sets of qualities are important: date palms were first selected, under these pedoclimatic conditions, for their dates, and the local selection of different date palms aims to meet a variety of uses and tastes, indeed provided by the diversity of *alkak, ta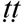agt* and *Ṣaεidi*, plus some others such as *a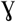zāl*. In other words, this PC1-PC2 space is the illustration of this local selection.

#### Cultivars, ethnovarieties and other categories

The variability within cultivar and category was assessed as the percentage of non-polymorphic loci between accessions from the same group. The five *Ṣaεidi* accessions (2398, 2399, 2400, 2401, 2404) appear 96.97% similar. Solely two samples over the five included (2400 and 2404) show a single different allele over the 17 loci. They cluster together in the NJ tree (Figure 2) and in the PCA plot (Figure 3ab).

The two accessions of *alkak* (2416 and 2051) cluster together in both PCA and NJ tree and are quite isolated from other accessions (Figures 2 and 3ab). They show 3 different alleles over 34 and are therefore very closely related but not perfect clones (91.18% identity). One of them, sampled in the first 2010 collection, was first labeled *fre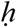i*, the local Arabic tradename of *alkak* for exportation to the Nile valley (see Battesti 2013: § 48, 90-91). This cultivar is maybe not a true cultivar, even though farmers do not think *alkak* can be obtained from seed. However, two samples are not enough to conclude. Further genetic analyses on more samples would help. Sample 2417 (*alkak wen žemb*) is closely related to these two accessions in both PCA and NJ tree (Figures 2 and 3ab). Interestingly, the farmers claim that *alkak* and *alkak wen žemb* are close relatives (wen *žemb* can be translated as “relegated”, “put aside”, i.e. a second class *alkak*). In contrast, *ta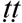agt* accessions (2052 & 2405) from two different collections are 100% similar meaning that this is probably a true cultivar although more samples are required to conclude. Identically, we will need further analyses to check the situation of *a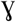zāl* (2409) and the very distant *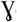rom a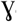zāl* (2065), which are locally supposed to be closely related cultivars — *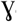rom* comes from *a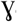ram* (pl. *i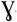arman*), the seed in Siwa language. Anyway, it would be a demonstrative argument of the ethnographic statement that is that “locally the shape does matter, but genes matter little” (Battesti 2013: § 62).

In the distance tree, the large clade comprises all feral accessions, with a few exceptions as some cultivated samples also group in this clade: *ta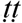agt* (2052 and 2405), *úšik ezzuwa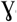* (2062, 2412, and 2058) and *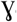rom a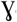zāl* (2065). In the PCA, the PC1 separates feral from cultivated samples with, again, the exception of *ta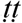agt* accessions that are distinguished from the desert feral group by PC3. Some geographical patterns seem at work for the feral date palms (Figures 2-3). The accessions 2426, 2427, and 2428 are part of a same clade in the distance tree, and the accession 2425 belongs to the next rank up: all four samples are from Taba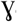ba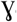, an oasis abandoned since Roman times. In the same clade are the samples 2414 from al-εAraj and 2432 from Timira, other nearby oases abandoned since Roman times. Accessions from the ancient oases Timira (2433, 2431, and 2432), Um Iktaben (2420 and 2413) and al-εAraj (2421, 2414) seem to present a higher genetic diversity. In ancient times, those locations were of course linked by desert trails. At that time, the main trail that linked the eastern extremity of Siwa region, i.e. εaīn Ṣafī, with Bahriyya oasis (and beyond with Giza and then Cairo) was going through al-εAraj.

The positions of *úšik* #1 and *úšik* #2 are interesting as our samples are scattered all over the distance tree (with a clear predominance of the garden seedlings *úšik* #1 within the clade of the garden named types date palms). In the PCA (Figure 2ab), PC1, PC2 and even PC3 do not offer a clear difference between the seedling palms kept in the gardens (*úšik* #1) and those that grow alone and without care in the palm grove outside the gardens (*úšik* #2). Nonetheless, should be noted that these *úšik* offer a real genetic diversity and they form a group overlapping the cultivated group and the desert feral group in the PCA.

### Morphometric analysis of seeds

The length and width of 1100 seeds from 55 date palms are presented in Figure 5. Feral date palms show significantly smaller seeds than cultivated date palms (Tukey’s test, p-value < 0.05) as previously demonstrated (Gros-Balthazard et al. 2016). Indeed, we expect the environment to play a major role in the size of the seeds. Cultivated date palms grow in a very favorable environment (fertilizer, water, care …) while feral date palms grow untended and thus produce smaller seeds.

**Figure 5.**
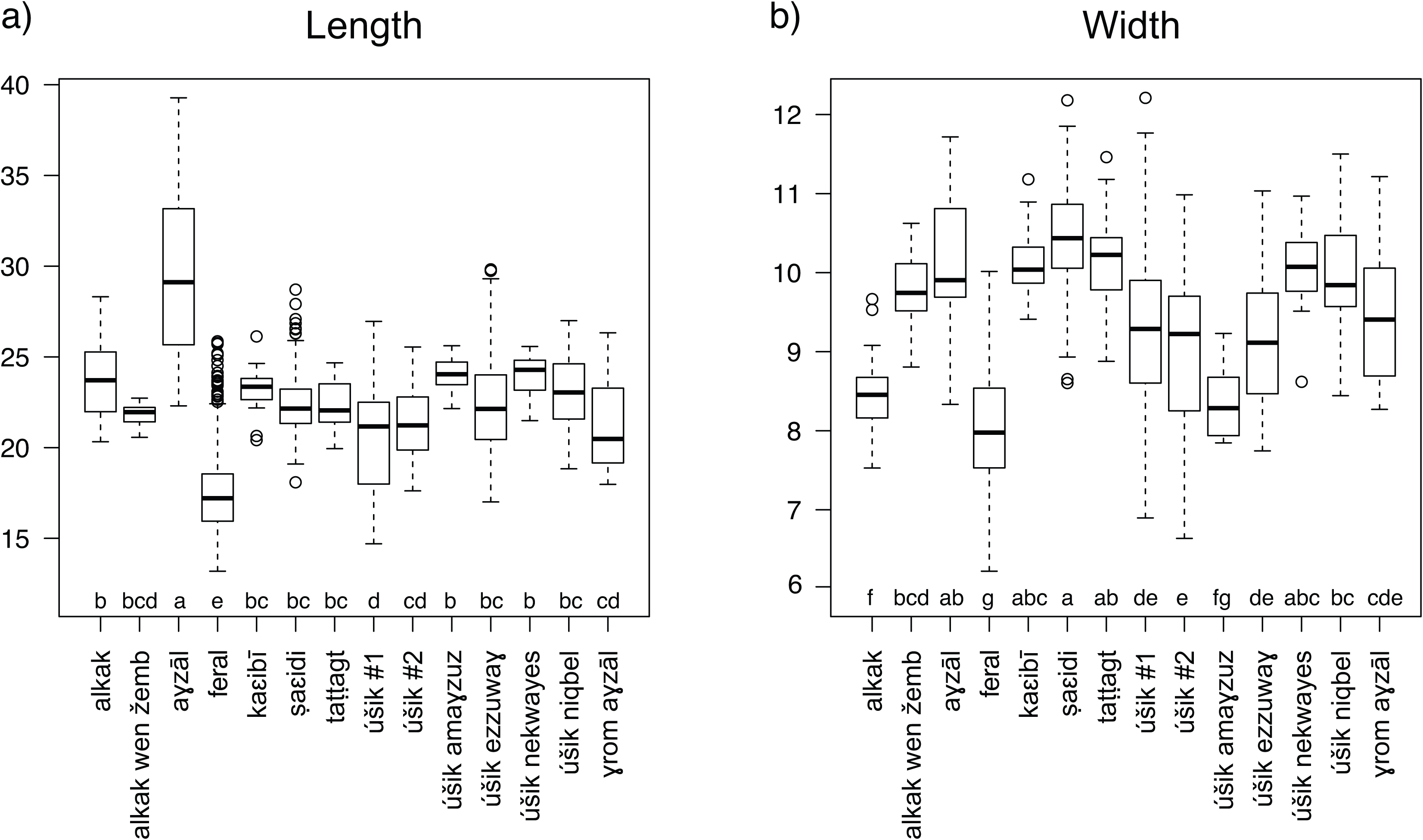
Seed size of Siwa date palms (mm). a) Length, b) Width. Letters below each box correspond to Tukey’s group.

**Figure 6:**
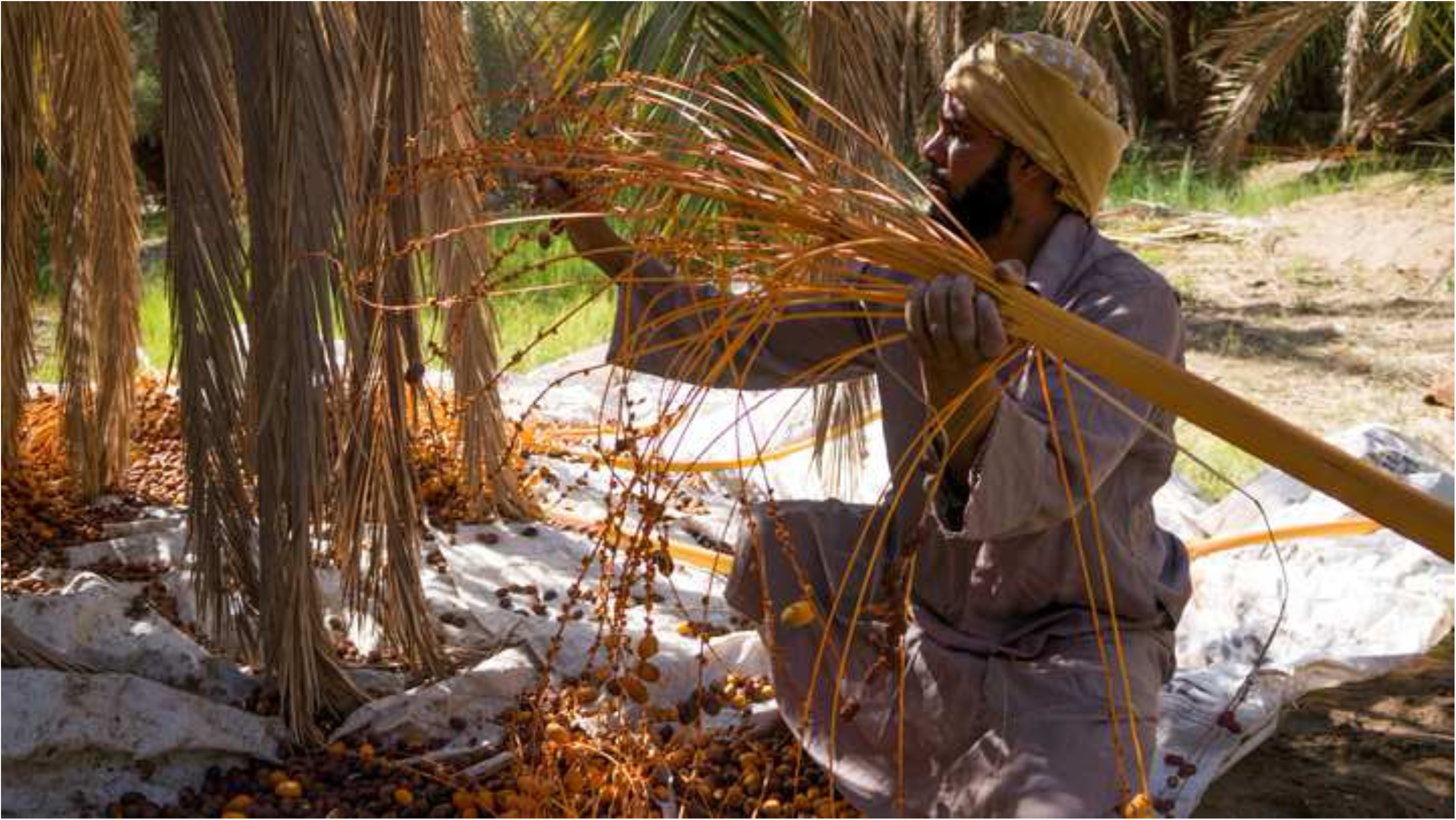
A farmer harvesting the dates of his garden in Abregani palm grove, Siwa oasis (Egypt), Nov. 15^th^, 2014. Vincent Battesti.

**Figure 7:**
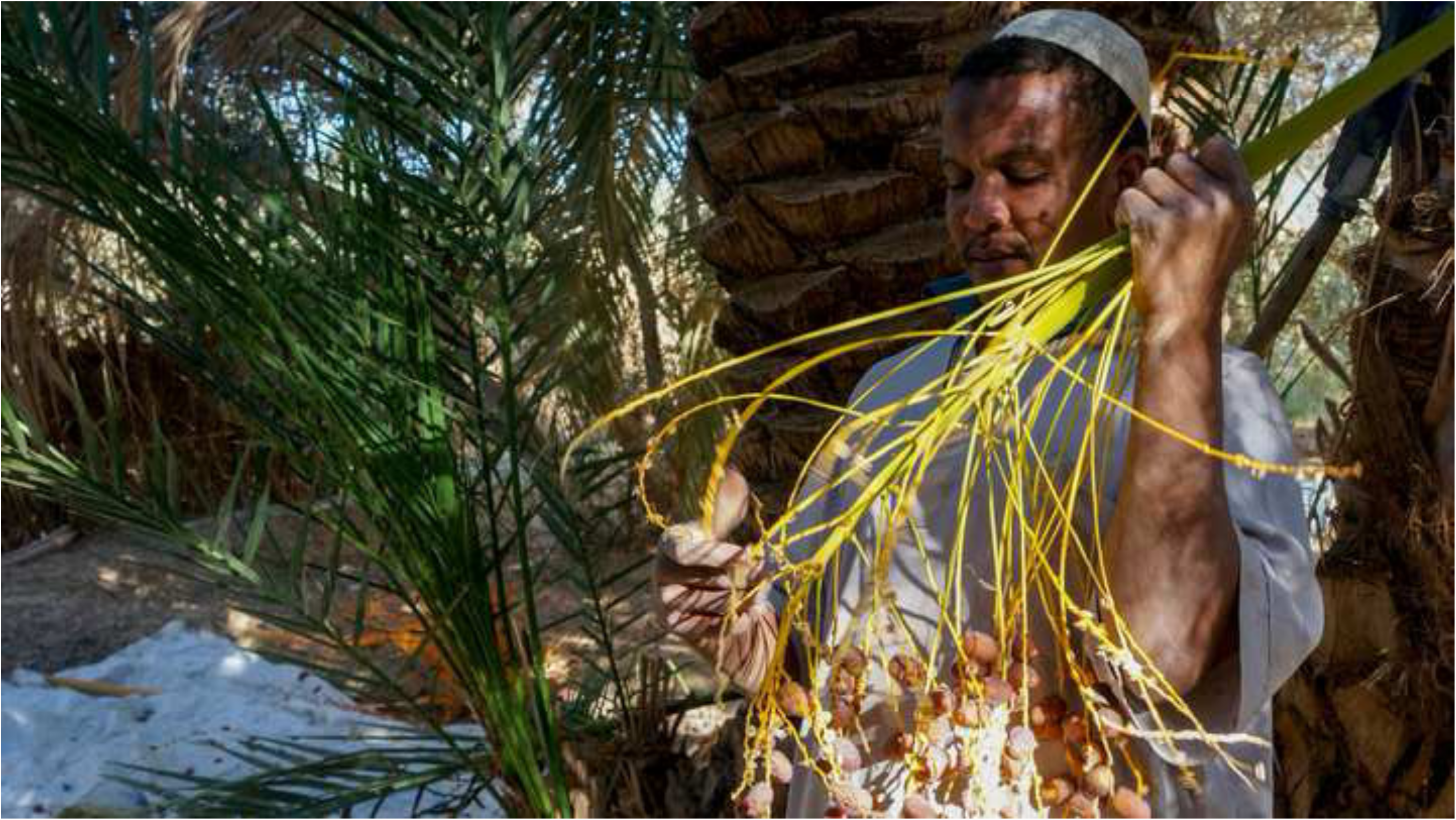
During the date harvest in a garden of Azzumuri palm grove in Siwa oasis (Egypt), Dec. 5^th^, 2013. Vincent Battesti.

**Figure 8:**
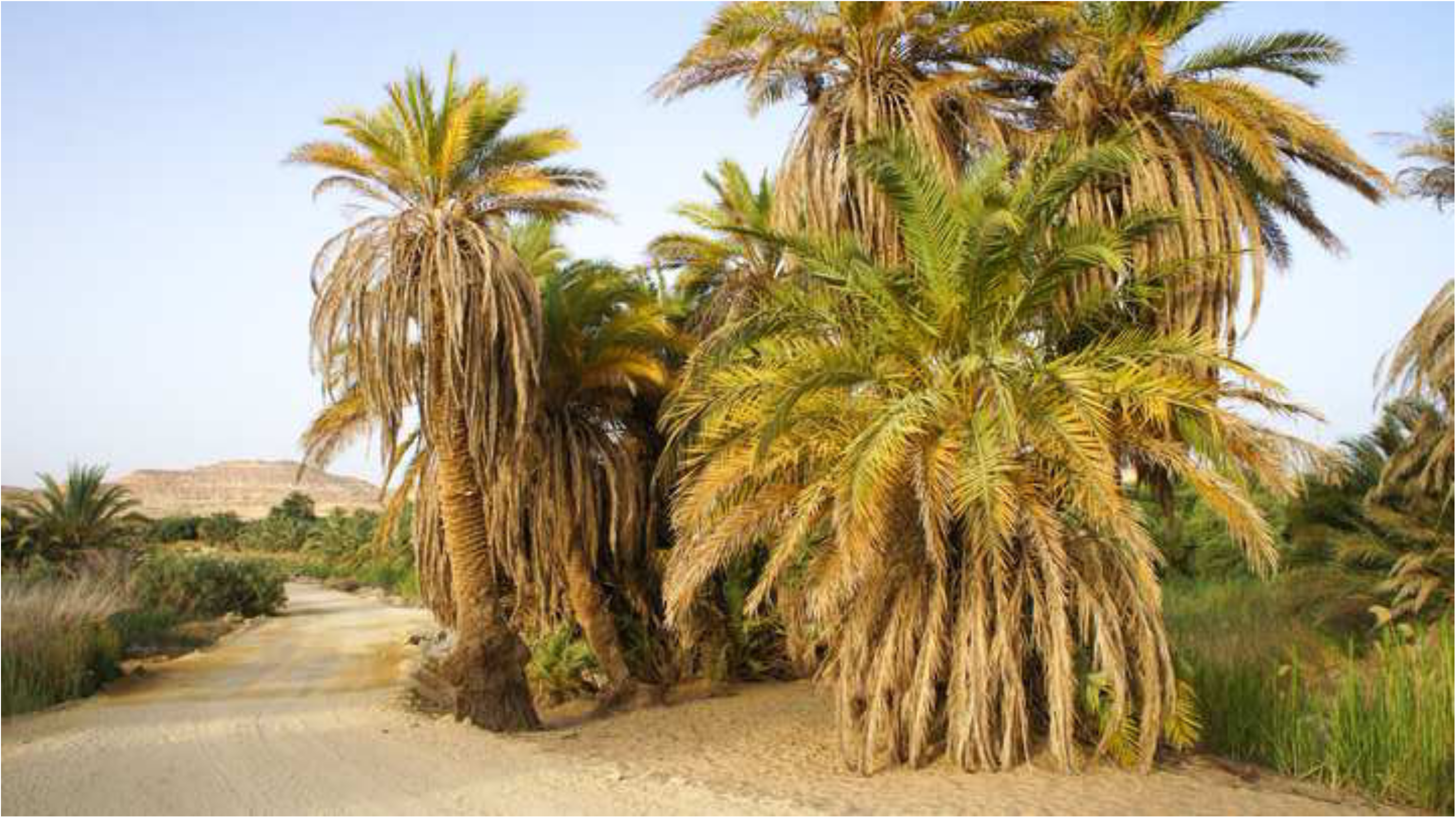
Untended seedling date palms, on the edge of the Baεed al-Dīn palm grove, West part of Siwa oasis (Egypt), May 21^st^, 2013, Vincent Battesti.

We estimated the variation of seed size and shape. Seeds from *úšik* #1, *úšik* #2 and desert feral have a very high variability, both in terms of size and shape (Table 4). This is in accordance with the fact that these grow in a diversity of conditions (regarding the size) and are not reproduced clonally (regarding the shape) but rather represent a group of unrelated samples. On the contrary, *Ṣaεidi* and *alkak*, which are supposedly groups of cloned accessions (cultivars), display lower seed shape and size variability (Table 4).

**Table 4.**
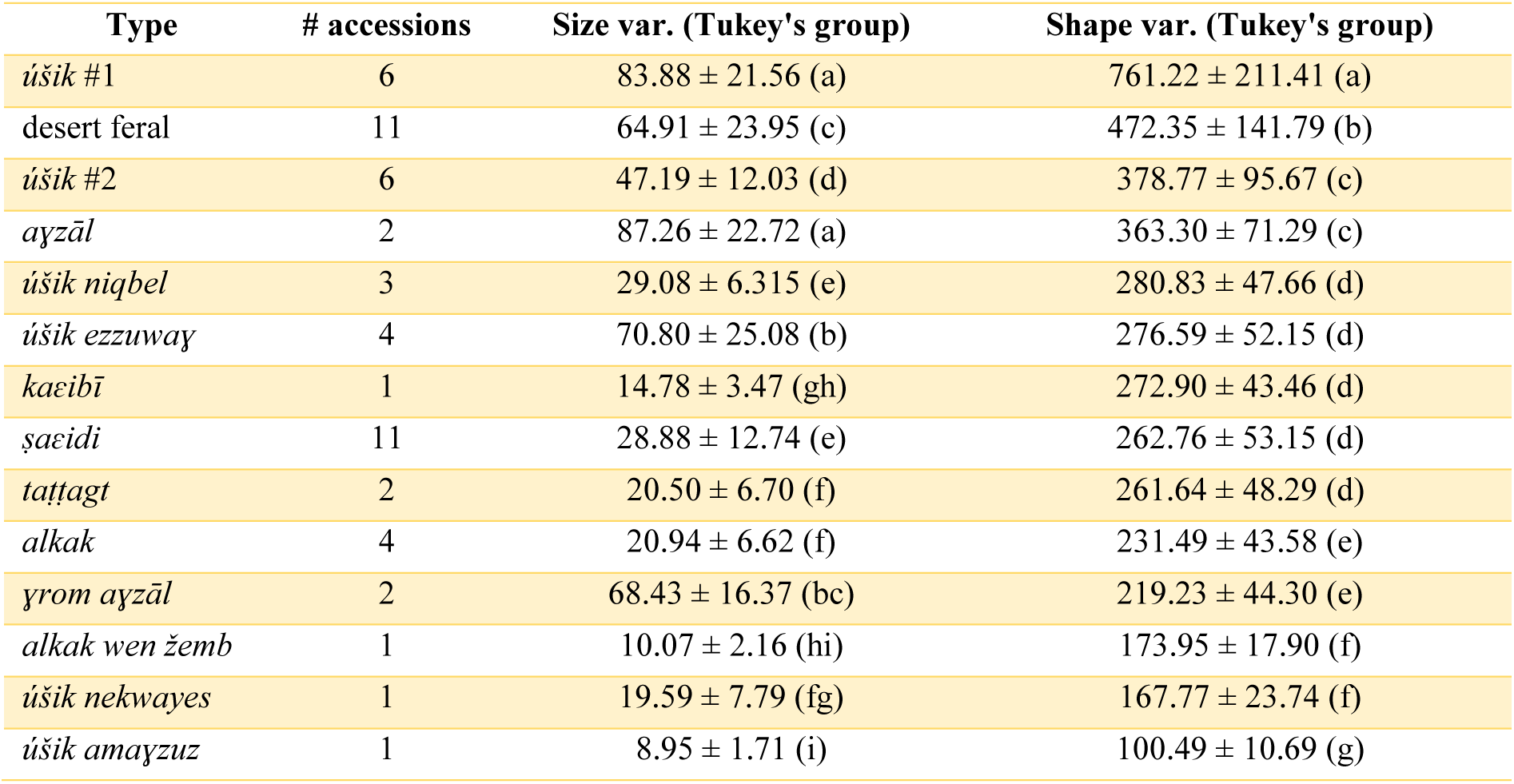
Variability of seed dimensions and shape within named types of Siwa. It is calculated as the dispersion of seeds around the related species’ centroid in two PCA spaces obtained from size parameters (Size Var.) and 64 Fourier coefficients related to dorsal and lateral seed shapes (Shape Var.) using the rarefaction method to balance sample size in each group. The values are the average over the mean distance between 10 randomly sampled seeds in one hundred replicates and the standard deviation over the one hundred replicates. The number of accessions available for each group is given (# accessions). The groups derived from Tukey’s test are given in parentheses.

| Type | # accessions | Size var. (Tukey's group) | Shape var. (Tukey's group) |
| --- | --- | --- | --- |
| *úšik* #1 | 6 | 83.88 ± 21.56 (a) | 761.22 ± 211.41 (a) |
| desert feral | 11 | 64.91 ± 23.95 (c) | 472.35 ± 141.79 (b) |
| *úšik* #2 | 6 | 47.19 ± 12.03 (d) | 378.77 ± 95.67 (c) |
| *ayzāl* | 2 | 87.26 ± 22.72 (a) | 363.30 ± 71.29 (c) |
| *úšik niqbel* | 3 | 29.08 ± 6.315 (e) | 280.83 ± 47.66 (d) |
| *úšik ezzuway* | 4 | 70.80 ± 25.08 (b) | 276.59 ± 52.15 (d) |
| *kaeibī* | 1 | 14.78 ± 3.47 (gh) | 272.90 ± 43.46 (d) |
| *šaēidi* | 11 | 28.88 ± 12.74 (e) | 262.76 ± 53.15 (d) |
| *taṭṭagt* | 2 | 20.50 ± 6.70 (f) | 261.64 ± 48.29 (d) |
| *alkak* | 4 | 20.94 ± 6.62 (f) | 231.49 ± 43.58 (e) |
| *γrom ayzāl* | 2 | 68.43 ± 16.37 (bc) | 219.23 ± 44.30 (e) |
| *alkak wen žemb* | 1 | 10.07 ± 2.16 (hi) | 173.95 ± 17.90 (f) |
| *úšik nekwayes* | 1 | 19.59 ± 7.79 (fg) | 167.77 ± 23.74 (f) |
| *úšik amayzuz* | 1 | 8.95 ± 1.71 (i) | 100.49 ± 10.69 (g) |

The PCA (Figure 4) does not speak a lot by itself besides proving quite clearly that seeds of desert feral and *úšik* have a higher variability than the garden named types as a whole. Nonetheless, we can guess that PC1 is mainly drawn by desert feral and *úšik* #1, PC2 by cultivated named types (and especially *Ṣaεidi*). PC3 is also mainly drawn by desert feral and *úšik* #1 and so the space PC2/PC3 slightly differentiates between *úšik* #1 (benefiting from care) and *úšik* #2 (outside the gardens) seedlings. Strangely enough, *úšik* #1 offers in PCA the same wide dispersal in seed shape, and even wider, than the desert feral date palms. As regards the “*úšik* xxx”, i.e. the date palms with a given name including the word *úšik*, like *úšik niqbel, úšik nekwayes, úšik ama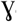zuz, úšik ezzuwa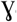*, their morphotypes roughly coincide with the other cultivated named types. They are locally known and said to have been former *úšik:* they are seedlings that became cultivars as they were selected from the local stock of *úšik* date palms and vegetatively reproduced.

**Figure 4.**
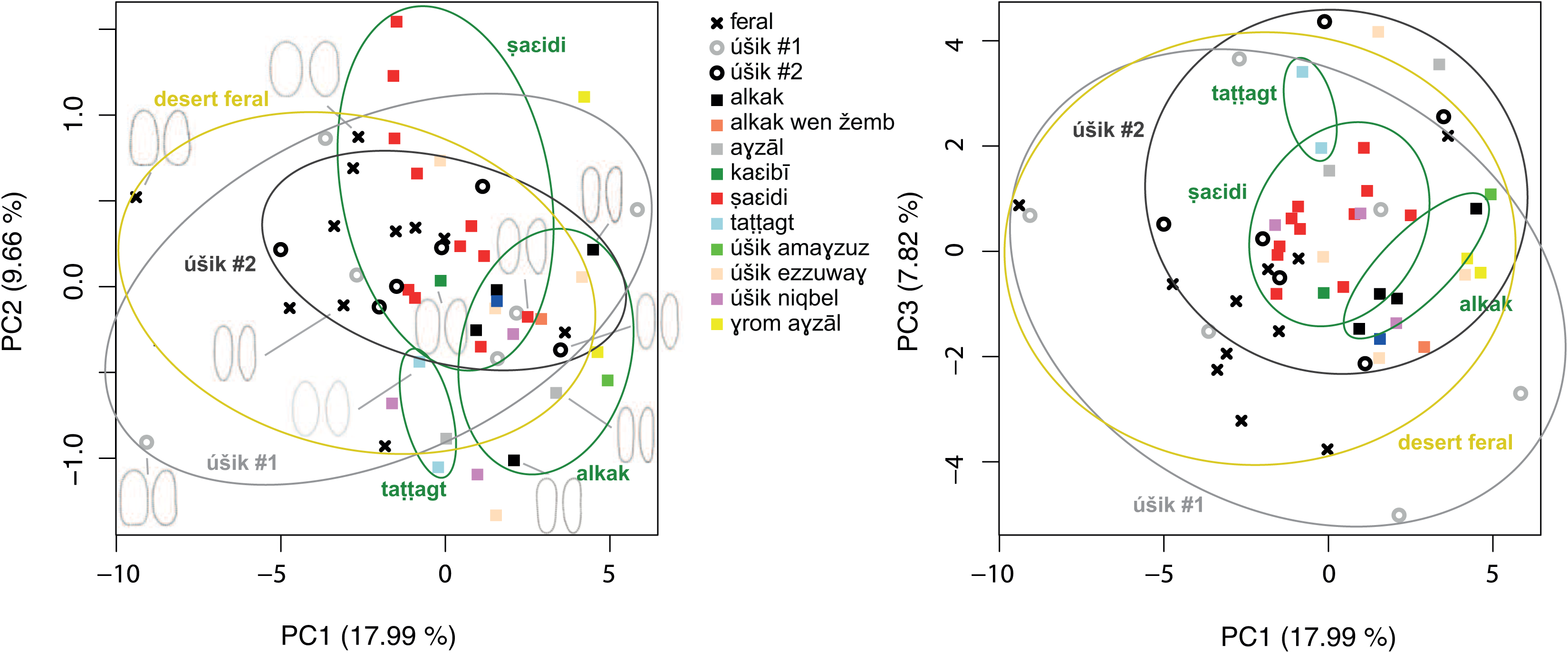
Principal Component Analysis (PCA) on 55 date palm accessions based on seed size and shape.

Using Linear Discriminant Analysis, we checked the probability of assigning date palms to their known status, uncultivated (desert feral and *úšik* #2, the palm grove seedlings) or cultivated (garden cultivated varieties and *úšik* #1, the garden seedlings). We found that 90.05% of cultivated and 79.21% of uncultivated date palms were assigned positively to their status meaning that these two groups show highly divergent seeds and that within each group, seeds display particular features. This supports again the distinction between these two groups (in the garden, out of the garden). A possible interpretation is that the uncultivated seeds have a richer palette of shape as a local genetic pool. We performed another LDA testing three groups: feral (desert feral) vs. named types (garden cultivated varieties) vs. seedlings (*úšik* #1 from the garden & *úšik* #2 from the palm grove). We found that 87.85% of named types, 78.82% of desert feral, and only 40.91% of garden and palm grove seedlings were assigned positively. In this case, it is interesting to note the inconsistency of the group of the *úšik: úšik* #1 are not *úšik* #2, all the seedlings do not form a consistent group and being adopted in the garden by a farmer makes a difference.

Identically, we performed assignation of seeds to the different cultivars (Table 5). We found that the 11 samples of *Ṣaεidi* and the 4 samples of *alkak* are easily assigned (respectively 80.38% and 79.72%), in accordance with the fact that the intra-varietal variability is low within *Ṣaεidi* and *alkak* samples and despite again the fact that those samples were collected in very different gardens of Siwa, so subjected to different cultivation conditions. The *úšik* #1 and #2 have a low cross-validation percentage in accordance with the fact that they are not clones. Surprisingly, the feral have a high CV even though they are not clones. This may be related to the fact that generation after generation, these not tended nor artificially selected palms show seeds that shift toward an “ideal” seed shape and size according to their environment through natural selection and the canalization process as explained in Siegal and Bergman (2002) and more specifically for date palms in Gros-Balthazard et al. (2016).

**Table 5.** Percent of cross validation (CV) for each variety/category.

| Type | # accessions | CV |
| --- | --- | --- |
| *úšik amayzuz* | 1 | 86.17 |
| *šaeidi* | 11 | 80.38 |
| *alkak* | 4 | 79.72 |
| desert feral | 11 | 79.04 |
| *úšik nekwayes* | 1 | 76.93 |
| *alkak wen žemb* | 1 | 73.38 |
| *taṭṭagt* | 2 | 73.11 |
| *γrom ayzāl* | 2 | 69.43 |
| *ayzāl* | 2 | 67.31 |
| *kaeibī* | 1 | 53.44 |
| *úšik #2* | 6 | 51.95 |
| *úšik #1* | 6 | 49.88 |
| *úšik ezzuway* | 4 | 48.19 |
| *úšik niqbel* | 3 | 43.37 |

## Discussion

In order to understand the intraspecific agrobiodiversity of a cultivated plant, it seems more than sensible to picture the local idea of this agrobiodiversity from the very people who make it exist, socially and biologically. Understanding this categorization of the date palm diversity was the objective of a former ethnobotanical article (Battesti 2013). To clarify the local categorization of this plant and its cultivars was one of the purposes of the present study and was realized using genetic markers. The second one was to check the reliability of the seed morphometrics tool: whether or not it gives congruent results with genetics.

The former ethnobotanical publication brought to light two facts: the first, quantitative, is that about fifteen or so named types (cultivars and ethnovarieties) exist in Siwa oasis; the second, qualitative, is that locally shape matters, but genes probably matter little. In other words, for the farmers, the resemblance produces identity, and this has implications on the richness of this resource. What seems interesting is to read biological sciences results annotated with local palm names. What does that mean? We make two systems dialogue: a scientific system based on genetic identity and a Siwa oasis farmers’ system based on phenotypic identity.

### Are cultivars of Siwa oasis vegetatively propagated (clones) or not?

The genetic analysis confirms the local Isiwan statement that the named types *Ṣaεidi* and *ta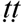agt* are true cultivars. The slight genetic difference between *Ṣaεidi* accessions (two samples over the five show a single different allele over the 17 loci) is possibly due to somatic mutation. The genetic analysis seems therefore to corroborate the idea that this locally named type is a cultivar: all the *Ṣaεidi* individuals belong to a single clone representing a true “cultivar”. They share not only a formal identity but also a genetic identity. Although these accessions were collected in distant gardens in all Siwa palm groves, they apparently all originate from the vegetative reproduction of a single ancestor. We were even able to correct a given name: the label first given during the collection for the accession 2400 was *úšik*; considering the result, we were able to double cross-check the data in the field and we get the confirmation that 2400 is locally named *Ṣaεidi* and was wrongly identified by the first informant as *úšik*.

Nonetheless, the genetic analysis should also have confirmed the local Isiwan statement that the named type *alkak* for instance is a true cultivar, but even very closely related, the two *alkak* accessions are not perfect clones. The ethnographic data strongly suggested us that *úšik nekwayes* and *kaεibī* are more likely each an ethnovariety (a collection of individuals with similar shapes, reproduced vegetatively) than each a true cultivar (a clone, originally a single individual reproduced vegetatively). In this study, we do not have the means to verify that.

Our study also suggests that the given names *úšik ezzuwa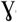* and *úšik niqbel* refer to ethnovarieties. Four samples from a single garden of Tamusi palm grove in Siwa illustrate the statement that “locally shape matters, but genes matter little”: two *úšik niqbel* (2423 & 2422) and two *úšik ezzuwa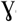* (2412 & 2419) are scattered in the NJ tree (Figure 2). These distributions are perturbing enough to think of a mistake made by the informant or of an operating error in the lab. Another hypothesis emerges and discards the idea of a mistake once we go back to the raw material: despite the genetic distance highlighted by the analysis, the dates of the 2422 *do look like* 2423 (fleshy, oblong and brown dates), which allows them to share the name of *úšik niqbel*, just as the dates of 2419 *do look like* 2412 (round and reddish/black dates), allowing them to share the name of *úšik ezzuwa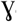* (“*ezzuwa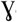*’ is related to the red color in Siwa Amazigh language). So the two samples of *úšik ezzuwa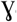* differ greatly from each other and ethnographic data partly predicted that as this given name had a “controversial status” (Figure S1) among farmers of Siwa oasis regarding the existence of a single clone. It seems that the answer, to be confirmed, is “no” as the four *úšik ezzuwa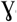* samples are scattered in the NJ tree. The case of *úšik niqbel* is more puzzling, as the two samples are also genetically different and this given name was supposed to be the name of a cultivar and not of an ethnovariety (Figure S1). The local primacy of form over gene is definitively corroborated by the vicinity in the NJ tree of the said *úšik ezzuwa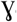* (2412) and the said *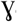rom a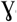zāl* (2065): they obviously share a common ancestor, but still, they are socially differentiated by different names because the genetic distance is there efficient enough to be apparent, for they are seen as displaying different forms (*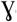rom a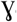zāl* bears yellow unripe dates and brown ripe dates).

Again, more data is necessary to check this, but named categories in Siwa could correspond to ethnovarieties as a general rule, with cultivars being the exceptions. That means that beyond the apparent agrobiodiversity (15 given names of cultivars), a “hidden genetic richness” exists (more than one clone per given name), all those clones duly selected, and reproduced by the local farmers. The ethnographic analysis combined with genetic analysis did confirm that the “genetic identity” — which is the classic and scientific definition of a “true” cultivar for the date palm — is not meaningful for local farmers; but “shape identity” makes sense for them. That means that a seedling date palm sharing similarity in shape (plant or fruit) with a known cultivar can be “added” to this cultivar and in a permanent way: it will eventually be vegetatively reproduced by farmers too and the cultivar will become an “ethnovariety”.

### Feral versus cultivated palms

Two facts emerge from this study regarding the feral status in contrast with the cultivated samples. The desert feral date palms could be the remains of a putative ancient gene pool from which were drawn a cultivar like *ta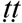agt*, but also some current *úšik* date palms (growing in the gardens or carelessly in the palm grove). Secondly, *úšik* date palms are a true gene pool for the farmers of the palm grove (in the past and present), offering a wide genetic and morphological diversity. They are a possible original pool from which stems part of the current diversity of cultivars. In this respect, *úšik* date palms — pl. *úškan*, maybe this local word can be compared to one of the two categories of feral in Jerid oases in Tunisia, *šeken* (Battesti 2005), with the same š and k consonants — should deserve a better attention in scientific analyses of agrobiodiversity. Fortunately, analyses of seed shape and size allow us to discriminate them, confirming the relevance of the archaeobotanical tool.

### Focusing on the fruits of date palms and their shape

For local farmers, the form of the date palm and its pedoclimatic requirements, and the form and quality of its fruit are the understandable keys to categorize their date palms agrobiodiversity. Maintaining such palm groves has as first objective to produce dates, with a diversity that meets export and local demands (different tastes, different preservation capacities, different harvest seasons). Locally, the form and quality of the fruits prevail over the genetic proximity, except that local farmers know that the best way to obtain another date palm bearing the required form and quality of fruit is to use vegetative reproduction: the best way, but not the only one. Our genetic and morphometric analyses are congruent but fail together sometimes to duplicate the local categorization of the date palm biodiversity, because of the local existence of ethnovarieties in Siwa. Is Siwa an exception or is it an overall and unnoticed pattern of date palm agriculture?

Finally, the ethnographic study (Battesti 2013) showed that even if the richness of the date palm heritage of Siwa is celebrated, and despite the apparent abundance of variety names in the literature, we have only about fifteen different given names. As said, an unknown portion of these given names are not “true” cultivars but “ethnovarieties” (the ethnographic study had already sorted out the issue of “categories” of local palms that could have been understood inaccurately as varieties of palms in the literature). The existence of ethnovarieties actually increases the genetic agrobiodiversity of the date palm in Siwa, hidden under a number of “given names”. Nonetheless, this number stays small in comparison with other oasis regions such as the Jerid in Tunisia where 260 varieties are recorded (Rhouma 1994, 2005, Battesti 2015). A hypothesis that possibly explains this low number is historical: when new communities established Siwa oasis — that may have been abandoned at that time — during the 11^th^ or 12^th^ century, they probably did so with a clear direction for an economy of exportation, and especially of two complementary elite cultivars: the *Ṣaεidi* for remote big cities and the *alkak* for the consumption of the Ṣaεidi-exporting people — the Bedouins, at least these last centuries (Battesti 2013). The genetic PCA space that explains the best the diversity of the samples’ genetic differences is manifestly a space stretched on its ordinate by the two elite (and exported) cultivars of Siwa (*Ṣaεidi* and *alkak*) and on its abscissa by the locally appreciated and consumed cultivars (but no exportable). So, how do we interpret that?

The proximity of all the feral accessions from ancient and abandoned oases in the same clade in the distance tree may suggest an old gene pool of cultivated date palms, which survives in nowadays cultivated date palms of Siwa through *ta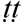agt* (2052 & 2405), *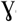rom a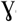zāl* (2065), some *úšik ezzuwa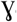* (2412) and some garden and palm grove subspontaneous date palms (*úšik* #1 and *úšik* #2). This hypothesis leads to its complementary: the other forms of date palm may have been imported to Siwa.

### Morphometries as a reliable tool to identify true cultivars?

This study validates the seed morphometric tool based on the analysis of seed size and shape; nonetheless that depends on the use we make of this tool. Morphometry is efficient with a “standard” or true cultivar, for instance here “*Ṣaεidi*”. If the morphometric PCA fails to regroup all the *Ṣaεidi* accessions, for instance, assignation of seeds through LDA is quite conclusive (Table 5): 80.38% of the 11 samples of *Ṣaεidi* were easily positively assigned. According to the ethnographical study, all the “Ṣaεidi” are locally said to be real clones — i.e. all the palms bearing the name *Ṣaεidi* are offshoots (vegetative reproduction) of “*Ṣaεidi*” palms, all originating from the same common ancestor (Battesti 2013). In this case, morphometric analysis clearly identifies all the samples bearing the name “*Ṣaεidi*” as having similar seed shape. Finally, the genetic analysis (with a Nei 1983 distances tree, NJ method) ascertains an obvious proximity of all *Ṣaεidi* samples we collected in Siwa oasis. But when archaeobotanists use this tool, they do it blind of any genetic results: the morphometric PCA differentiates clearly between *ta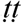agt* and *alkak*, for instance, but it will be more difficult to distinguish between *Ṣaεidi* and *alkak*, as they overlap maybe too significantly. Thus, the morphometric analyses will not always ascertain archaeobotanists that such a seed belongs to such a named type. However, it is rarely the aim with archaeological material. This tool, on the other hand, should offer the possibility to work on a collection of archeological seeds to check how they are scattered on the PCA space, and to deduce 1) that they belong to differentiated clones, 2) the presence of feral versus cultivated date palms.

## Conclusion

We would like to underline the originality of the study, involving three different methodological and disciplinary approaches. Each discipline still has its own agenda, and this has to be kept in mind all along the process of cooperation, from the collection of the data to their interpretation to satisfy the expectations of all. This is not only for the satisfaction of each of the participants, but to offer the opportunity for all to cooperate at their best: for instance, to maintain the traceability of each sample, not only at the level of the region, or cultivated/uncultivated, but more precisely at the very level of the garden (or GPS location) or even the identity of the informant, because these data can provide socioecological information, especially to assess the quality of the data. The date palm agrobiodiversity is the result of and is only due to the work of generations of the local farmers. So, it is not only a matter of observing the good practices: good scientific practice cannot simply ignore local knowledge, and this requires a social science expertise that cannot be invented or cannot be improvised in the field: during the collection of samples, the local categorization that makes sense for local farmers has to be understood by the collector. The terms of a disciplinary dialogue are important to define because cooperation will necessarily continue. A forthcoming publication will analyze an expanded collection of genetic samples of Siwa, which should enable us to better answer this question: are local varieties of Siwa cultivars or ethnovarieties? And this forthcoming study will possibly help us to better understand the biogeography of the Siwa date palms.

## Captions for illustrations

The date palm (*Phoenix dactylifera* L.) in Siwa oasis (Egypt): How ethnographic, morphometric, and genetic analyses together explain the local agrobiodiversity Vincent Battesti, Muriel Gros-Balthazard, Clémence Ogéron, Sarah Ivorra, Jean-Frédéric Terral, Claire Newton

## Supporting Information

**Figure S1.**
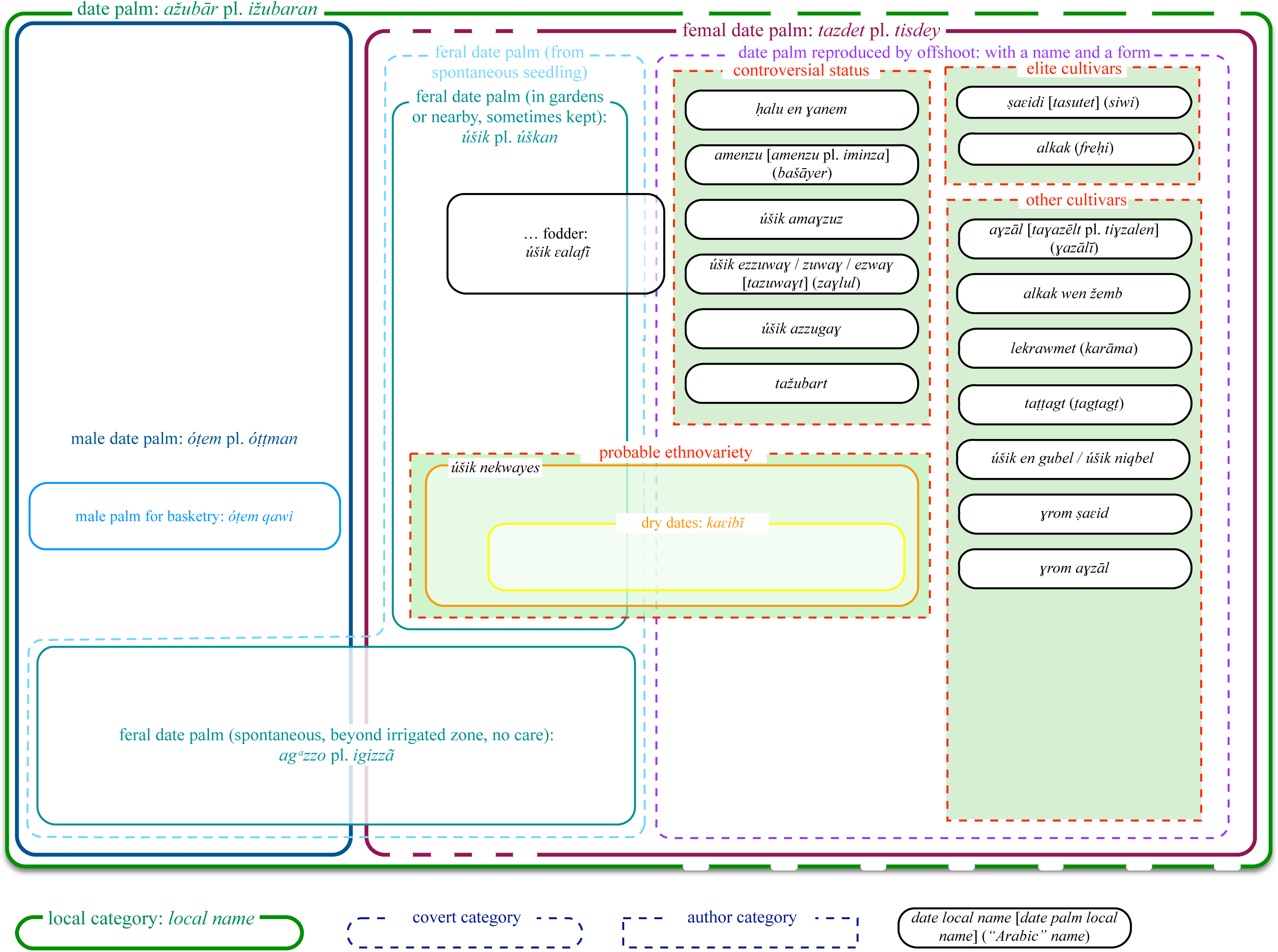
Date Palms Categorization in Siwa oasis

